# Genome-scale metabolic models reveal determinants of phenotypic differences in non-Saccharomyces yeasts

**DOI:** 10.1101/2023.05.09.539830

**Authors:** Jakob P. Pettersen, Sandra Castillo, Paula Jouhten, Eivind Almaas

## Abstract

**Background:** Use of alternative non-*Saccharomyces* yeasts in wine and beer brewing has gained more attention the recent years. This is both due to the desire to obtain a wider variety of flavours in the product and to reduce the final alcohol content. Given the metabolic differences between the yeast species, we wanted to account for some of the differences by using *in silico* models.

**Results:** We created and studied genome-scale metabolic models of five different non-*Saccharomyces* species using an automated processes. These were: *Metschnikowia pulcherrima, Lachancea thermotolerans, Hanseniaspora osmophila* and *Kluyveromyces lactis*. Using the models, we predicted that *M. pulcherrima*, when compared to the other species, conducts more respiration and thus produces less fermentation products, a finding which agrees with experimental data. Complex I of the electron transport chain was to be present in *M. pulcherrima*, but absent in the others. The predicted importance of Complex I was diminished when we incorporated constraints on the amount of enzymatic protein, as this shifts the metabolism towards fermentation.

**Conclusions:** Our results suggest that Complex I in the electron transport chain is a key differentiator between *Metschnikowia pulcherrima* and the other yeasts considered. Yet, more annotations and experimental data have the potential to improve model quality in order to increase fidelity and confidence in these results. Further experiments should be conducted to confirm the *in vivo* effect of Complex I in *M. pulcherrima* and its respiratory metabolism.

## Background

In recent years, there has been increased interest in using alternative non-*Saccharomyces* yeasts for beer and wine brewing[1, 2, 3, 4, 5, 6, 7]. In general, there are two primary drivers for the adoption of non-*Saccharomyces* fermentation strains: First, wine producers aim to decrease the resultant alcohol content in their products. Second, some brewers seek to enhance the complexity of aroma compounds, thereby emulating the rich flavor profile of spontaneously fermented beverages. In this study, our primary focus will be on the pursuit of reduced alcohol content.

Climate change has resulted in warmer and sunnier summers in wine producing regions, leading to higher sugar content in ripe grapes. When the must of high sugar grapes are fermented, this leads to higher alcohol content in the product. As a consequence, the alcohol content of wine has risen by approximately 1% alcohol by volume each decade since the 1980s in some wine producing regions[8, 9]. Whereas approaches such as dilution of the must, earlier harvesting of the grapes, and post-fermentation removal of alcohol can bring down the resulting alcohol content, such approaches come at the expense of diminished oenological qualities as well as breaking with established standards for wine brewing[10]. Additionally, these practices may violate local, national, and international regulations, such as the OIV Codex[11, 12]

In order to create wines with reduced alcohol content without losing the rich flavours, aeration during the fermentation process has been proposed as a solution[6]. Unfortunately, using this approach with the canonical wine yeast *Saccharomyces cerevisiae* has proven to be challenging. First of all, the most common strains of *S. cerevisiae* are Crabtree positive, meaning that glucose predominantly gets fermented to ethanol even when oxygen is available[13, 14, 15]. Furthermore, aeration often leads to the production of acetic acid, which is considered an undesired by-product[15, 16]. On the other hand, many non-*Saccharomyces* yeasts produce less acetate and are Crabtree negative[17, 18, 2].

Using non-*Saccharomyces* yeasts alone is usually not a good option due to production of bad-tasting compounds and their low tolerance to ethanol. The latter short-coming leads to stuck fermentations and poor wine quality[16, 6]. Experiments attempting simultaneous inoculations of *S. cerevisiae* and non-*Saccharomyces* strains have revealed that exposure of *S. cerevisiae* to oxygen causes unacceptable amounts of acetate production, even after aeration is turned off[16]. In order to mitigate this problem, a technique with sequential inoculation has been developed. In this method, the must is inoculated with the non-*Saccharomyces* yeast with air sparging for two to three days before *S. cerevisiae* is added in order to complete the fermentation. This has proven to be a more viable approach for production of wine with reduced alcohol content, as the production of acetic acid remains low[19, 20, 21, 22].

In order to explain and predict such metabolic properties of yeast, genome-scale metabolic models (GEMs) have become a widely used tool[23, 24, 25, 26, 27, 28, 29]. For the model organism *S. cerevisiae*, well curated models exist [30, 31] which have been used for a variety of purposes. One application is for the explanation of the Crabtree effect using enzyme constrained genome scale models (ecGEMs)[26, 25, 32]. The ecGEMs incorporate enzymes’ turnover numbers and masses for constraining the internal metabolic fluxes, as the total mass which can be allocated for enzymatic proteins is limited.

In contrast to *S. cerevisiae*, GEMs are not readily available for most non-*Saccharomyces* yeasts. Nevertheless, the development of tools that enable automatic generation of these models from genomic data presents a possible solution to address this limitation. A promising approach, known as “carving” as described by Machado and coworkers[33], involves generating models from a meticulously curated universal model that serves as a comprehensive database of interconnected biochemical reactions. Furthermore, tools exist also for the incorporation of enzymatic constraints [34, 35] by automatically querying databases for protein masses and turnover numbers in order to integrate these data into an ecGEM.

In this article, we constructed GEMs for five of the most commonly applied non-*Saccharomyces* yeast strains attempted in wine brewing[4, 36, 7]. These are: *Hanseniaspora osmophila, Kluveromyces lactis, Metschnikowia pulcherrima, Torulaspora delbrueckii*, and *Lachancea thermotolerans*. The models were automatically constructed from genome data and carved form a curated universal yeast model. Using the reconstructed GEMs, we predict physiological properties of the yeasts *in silico*.

## Results

### Complex I differentiates *Metschnikowia pulcherrima* from the other yeast species

GEMs of the five non-*Saccharomyces* yeast strains were created by using CarveFungi[37]. From these models, protein constraints were incorporated, and ecGEMs (sMO-MENT) were made with AutoPACMEN[34] (see Methods for details). Key properties of the models are summarized in Table 1. We begin our investigation of the models’ phenotypic properties by predicting batch culture growth using dynamic FBA (dFBA)[38] simulations for 12 hours. We first use models without enzymatic constraints (Figure 1) and include the *S. cerevisiae* model iND750[39] as a reference. We choose glucose as the sole carbon source, with the initial concentration set to 10 mmol L^−1^ (1.8 g L^−1^). The supply of oxygen was restricted to 10 mmol*/*g DW Biomass*/*h (corresponds to 180 mg*/*g WD */*h). See Methods for uptake kinetics and additional details on the simulations.

**Table 1.** Properties of the GEMs studied in this paper. sMOMENT versions of the models have more reactions than the listed numbers, as autoPACMEN adds auxiliary reactions and splits reactions drawing from the protein pool into separate forward and backward reactions. The model for *S. cerevisiae* was taken from an external source[39] and did not have any corresponding enzyme constrained model.

| Organism | Reactions | Reversible reactions | Metabolites | Reactions drawing from protein pool |
| --- | --- | --- | --- | --- |
| *Metschnikowia pulcherrima* | 2049 | 610 | 1633 | 1310 |
| *Lachancea thermotolerans* | 2049 | 618 | 1647 | 1319 |
| *Torulaspora delbrueckii* | 1876 | 559 | 1510 | 1163 |
| *Kluyveromyces lactis* | 2131 | 621 | 1774 | 1401 |
| *Hanseniaspora osmophila* | 1556 | 520 | 1218 | 902 |
| *Saccharomyces cerevisiae* | 1266 | 436 | 1059 | NA |

**Figure 1.**
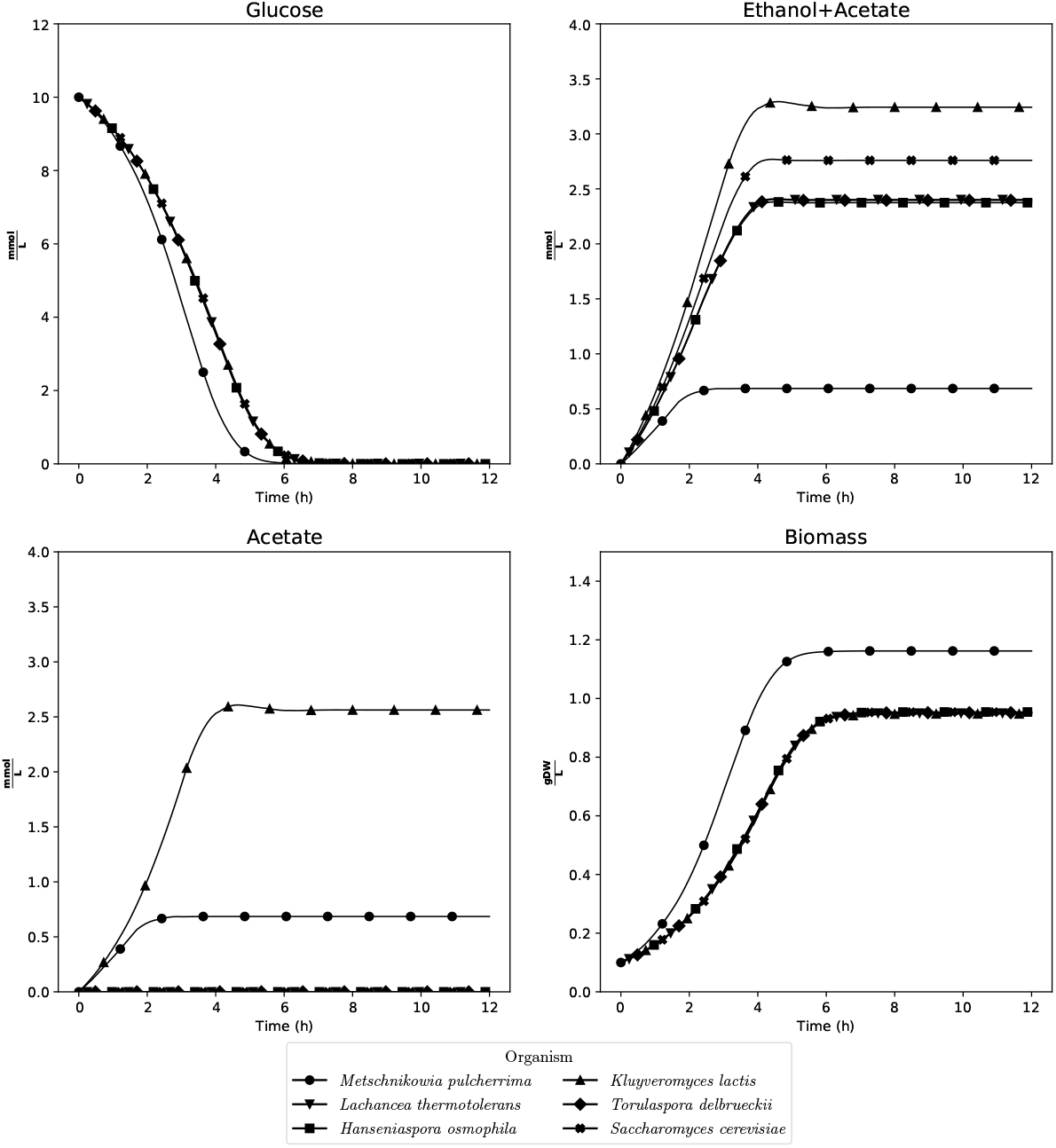
dFBA simulations of the models without enzymatic constraints for the six yeast models, starting with 10 mmol L^−1^ glucose. 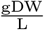: Grams of dry weight per liter.

We observed that simulation results for *M. pulcherrima* are quite different from the other yeasts, since less fermentation (production of ethanol and acetate) was undertaken compared to the other yeasts, and the growth dynamics resulted in higher biomass yield. We also initiated the simulations with 1000 mmol L^−1^ (180 g L^−1^) glucose, which is a more realistic sugar concentration in grape must used in wine fermentation (Supplementary Figure S1). This resulted in a higher degree of fermentation due to the fact that the balance between glucose and oxygen availability was shifted. Still, the same tendencies of *M. pulcherrima* to produce less fermentation products and attain higher biomass, were evident.

Subsequently, our objective was to elucidate the underlying reasons for the distinct metabolic phenotypes exhibited by *M. pulcherrima* compared to *Kluyveromyces lactis* and other yeast strains. To address this, we conducted a comparative analysis between the metabolic models of *M. pulcherrima* and *K. lactis*, excluding enzymatic constraints from consideration. This approach aimed to identify possible causal factors stemming from variations in metabolic network connectivity.

Directly comparing their reaction content, we found that 191 of the reactions in the *M. pulcherrima* model were not present in the *K. lactis* model. To probe functional consequence of these reactions, we sequentially (and cumulatively) removed each of these *M. pulcherrima* reactions and optimized for biomass production. The nutrient environment used for this assessment was identical to the one used for initiating the dFBA simulations. Some of the reactions were essential and therefore reinserted into the model before continuing. Of the considered reactions which were not essential, we observed two reactions which altered the growth rate: Complex I in the respiratory electron transport chain (NADH dehydrogenase), and mitochondrial Methylenetetrahydrofolate dehydrogenase (NAD+). Note that, removal of Complex I alone was sufficient to produce the same growth as in *K. lactis*. Conversely, adding the Complex I reaction to the *K. lactis* model yielded the same growth rate as of *M. pulcherrima*. Not surprisingly, removal of mitochondrial Methylenetetrahydrofolate dehydrogenase (NAD+) from *M. pulcherrima* did not have any effect on its own, nor did addition of the same reaction into the model of *K. lactis*: Since Complex I pumps protons and the mitochondrial Methylenetetrahydrofolate dehydrogenase does not, Complex I is beneficial when growth is APT dependent. Therefore, we chose to focus on Complex I when further comparing the models.

According to the reconstructions, *M. pulcherrima* was annotated with Complex I, whereas none of the other yeast strains contain this reaction. *Saccharomyces, Kluveromyces, Torulaspora, Lachancea*, and many other yeasts do not have the canonical Complex I of the electron transport chain, but instead feature an alternative Type II NADH dehydrogenases which does not pump protons across the mitochondrial membrane[40, 41, 42]. According to the GEM, Complex I pumps 4 protons across the mitochondrial membrane for each molecule of NADH being reduced, whereas the alternative Type II NADH dehydrogenases do not possess this ability. Hence, *M. pulcherrima* is able to create a larger proton-motive force (PMF) per mole of NADH being oxidized, which in turn increases the efficiency in generation of ATP per mole of glucose.

In order to obtain further evidence that Complex I was indeed present in *M. pulcherrima*, we conducted a BLAST[43] search with the protein sequences of *M. pulcherrima* against proteins annotated with Complex I functionality (EC number 7.1.1.2). This search was conducted as a blastp search through UniProt’s web portal[44] using standard settings. Our query returned matches to three manually curated Complex I subunits in Swiss-Prot[44], all for *Neurospora crassa* with evidence on transcript level: NADH-ubiquinone oxidoreductase 19.3 kDa subunit, mitochondrial; NADH-ubiquinone oxidoreductase 23 kDa subunit, mitochondrial; and NADH-ubiquinone oxidoreductase 24 kDa subunit, mitochondrial. The similarity to these sequences were (with corresponding *E*-value): 82% (2.8 · 10^−106^), 73.4% (1.6 · 10^−109^), and 53.9% (1.5 · 10^−84^), respectively.

In light of these discoveries, we suspected that by removing the advantage of proton pumping in Complex I, the metabolism of *M. pulcherrima* would become more similar to that of the other yeast strains. We therefore artificially changed the stoichiometry of the reaction to two or zero protons being pumped for each molecule of NADH consumed. We conducted a new set of dFBA simulations without enzyme constraints and with the same starting conditions as earlier, using *K. lactis* (lacking Complex I) as a baseline(Figure 2). From these results, we observe that the glucose consumption and biomass production are more or less identical for *M. pulcherrima* and *K. lactis* when the proton pumping is turned off. These results were identical to that of knocking out Complex I completely. Additionally, in the case of the partially inhibited state where two protons are pumped, the biomass yield and glucose consumption exhibit intermediary values, positioned between those observed in the wild-type and fully inhibited states. For the production of ethanol and acetate, it was observed that the production of ethanol and acetate decreased with the number of protons pumped by Complex I, yet *K. lactis* still displayed a higher production of ethanol and acetate when no protons were pumped.

**Figure 2.**
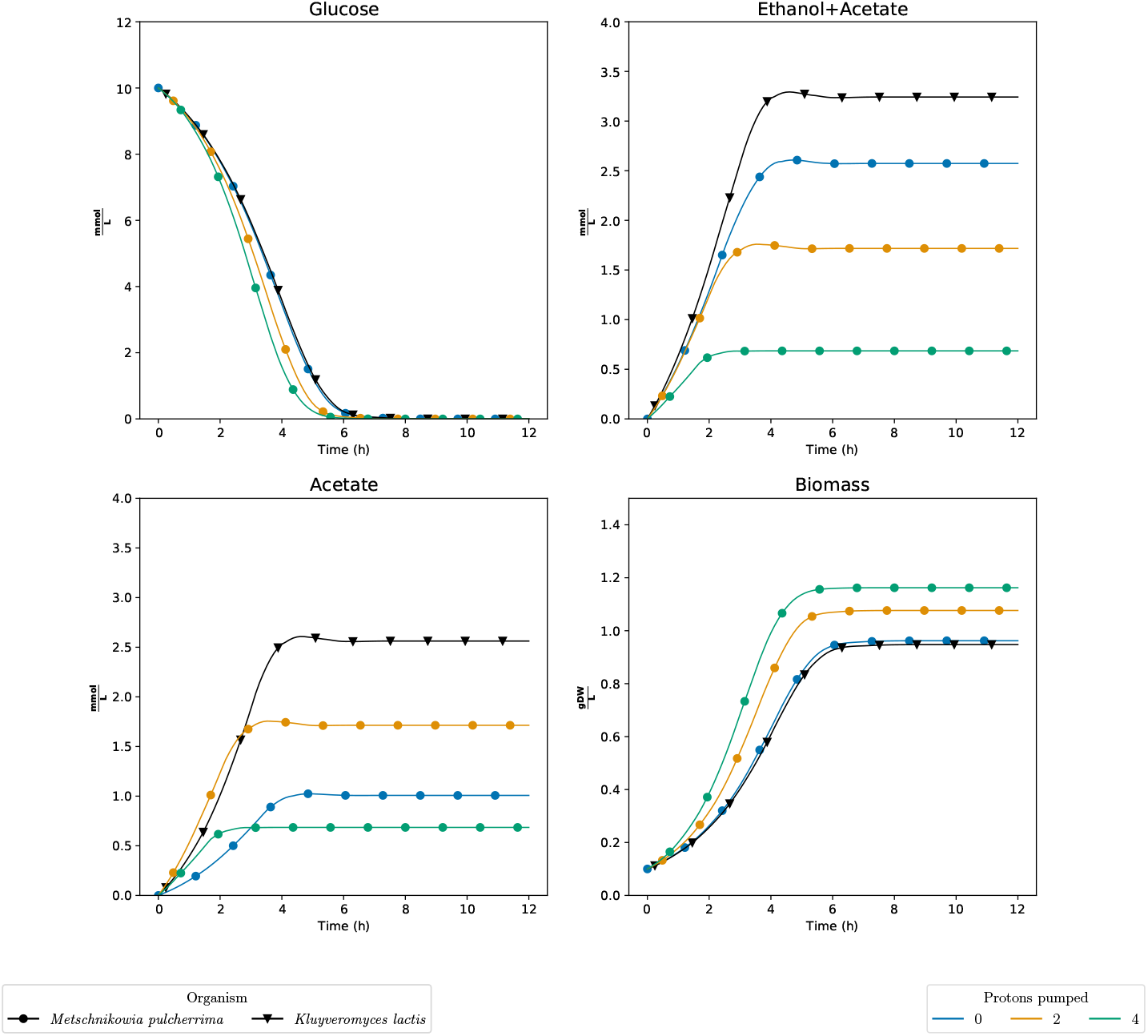
dFBA simulations of the models without enzymatic constraints for *Metschnikowia pulcherrima* and *Kluyveromyces lactis* when artificially changing the stoichiometry of the number of protons pumped by Complex I.

### Two reactions explain reduced production of fermentation products from *M. pulcherrima* in absence of Complex I activity

*K. lactis* exhibited a greater production of fermentation byproducts (acetate and ethanol) compared to *M. pulcherrima* even in the presence of inactive Complex I. This observation suggested the involvement of supplementary reactions. To explore this hypothesis, we deactivated Complex I in *M. pulcherrima* and performed cumulative knockouts of the reactions exclusive to this organism. We then assessed the combined production of acetate and ethanol following each knockout event to elucidate the potential role of these unique reactions in the observed differences. By this strategy, we found two reactions accounting for the difference in fermentation products. These two reactions were L-glutamate:NADP+ oxidoreductase (Equation 1) and Isocitrate:NADP+ oxidoreductase (Equation 2), running in the directions illustrated by the equations:

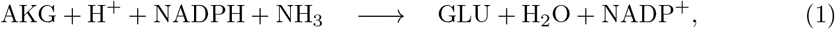

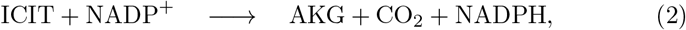

where AKG is *α*-ketoglutarate, GLU is L-glutamate, and ICIT is Isocitrate. The reactions are complementary, meaning that both reactions had to be knocked out in order to observe an increase in fermentation products (sum of ethanol and acetate). Conversely, adding either of these two reactions to the model of *K. lactis* resulted in a decrease in fermentation products. This complementary nature of the reactions is strange given that the two reactions constitute two consecutive steps in a pathway. We also conducted dFBA simulations where these two reactions in *M. pulcherrima* were knocked out (Supplementary Figure S2). We found growth, glucose consumption, and total fermentation to be almost identical in the two models when Complex I was left inoperative, whereas there were no major changes when Complex I was active.

### Protein constraints result in changes in use of metabolic pathways

Considering that Complex I is a key differentiator for *M. pulcherrima* with infinite amounts of enzymatic protein available, we next studied how the activity of Complex I affected metabolism when the available enzyme pool was constrained.

When simulating the effect of Complex I stoichiometry of *M. pulcherrima* with dynamic enzyme constrained FBA (decFBA)[26, 45] (Figure 3), we could not observe any major effect of the stoichiometry of Complex I for low availability of enzymatic protein. However, at the highest chosen protein pool level, the biomass yield was higher the more protons were pumped by Complex I. This is as expected, since decFBA becomes equivalent to dFBA when the available enzyme pool approaches infinity. For intermediate levels of the protein pool, the effects of stoichiometry were marginal. Most likely, this means that other metabolic pathways are chosen when availability of enzymatic protein is scarce, and hence, is not reliant on Complex I to the same degree. Moreover, the production of acetate and ethanol was affected by the number of protons pumped at the highest level of the enzyme pool only.

**Figure 3.**
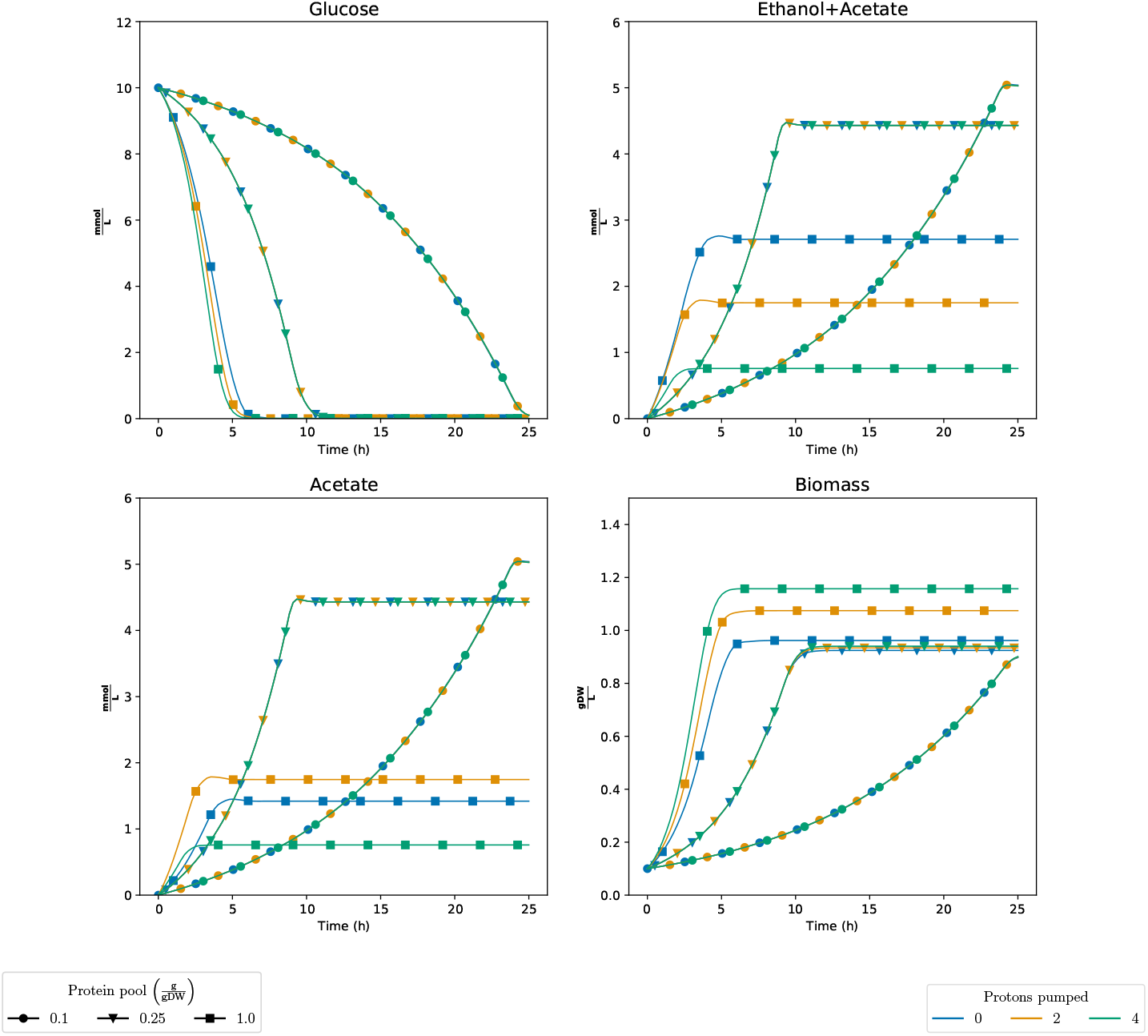
decFBA simulations of the sMOMENT model of *Metschnikowia pulcherrima* when artificially changing the stoichiometry of the number of protons pumped by Complex I under different levels of the protein pool. 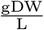 Grams of dry weight per liter.

Finally, we compared the sMOMENT models of *M. pulcherrima* and *K. lactis* as to get an overview of how the species compare when the access to enzymatic protein was restricted (Figure 4). As expected, the growth rate increased with the protein availability, but only up to a certain point where the substrate uptake rates became limiting. Also, the results show that the growth rate and biomass yield for *M. pulcherrima* was higher than for *K. lactis* at high availability of enzymatic protein. For lower levels of the enzyme pool, the two strains grew almost equally fast until glucose was exhausted. However, the biomass yield was marginally higher for *M. pulcherrima* than for *K. lactis*, even at the low level of protein, an observation which is due to differences in *k*_*cat*_ values in the two models. In addition, *M. pulcherrima* produced less acetate than *K. lactis* for all levels of the enzyme pool. Respiration is energetically more efficient than fermentation in utilization of the carbon source, but comes with a higher protein cost per unit of ATP produced[32, 25]. For this reason, we would expect to observe fermentation at high levels of enzymatic protein, in agreement with our observations. For the lowest level of enzymatic protein, Complex I is less relevant, but the difference in fermentation products can still be explained by the presence of L-glutamate:NADP+ oxidoreductase and Isocitrate:NADP+ oxidoreductase as we discussed earlier.

**Figure 4.**
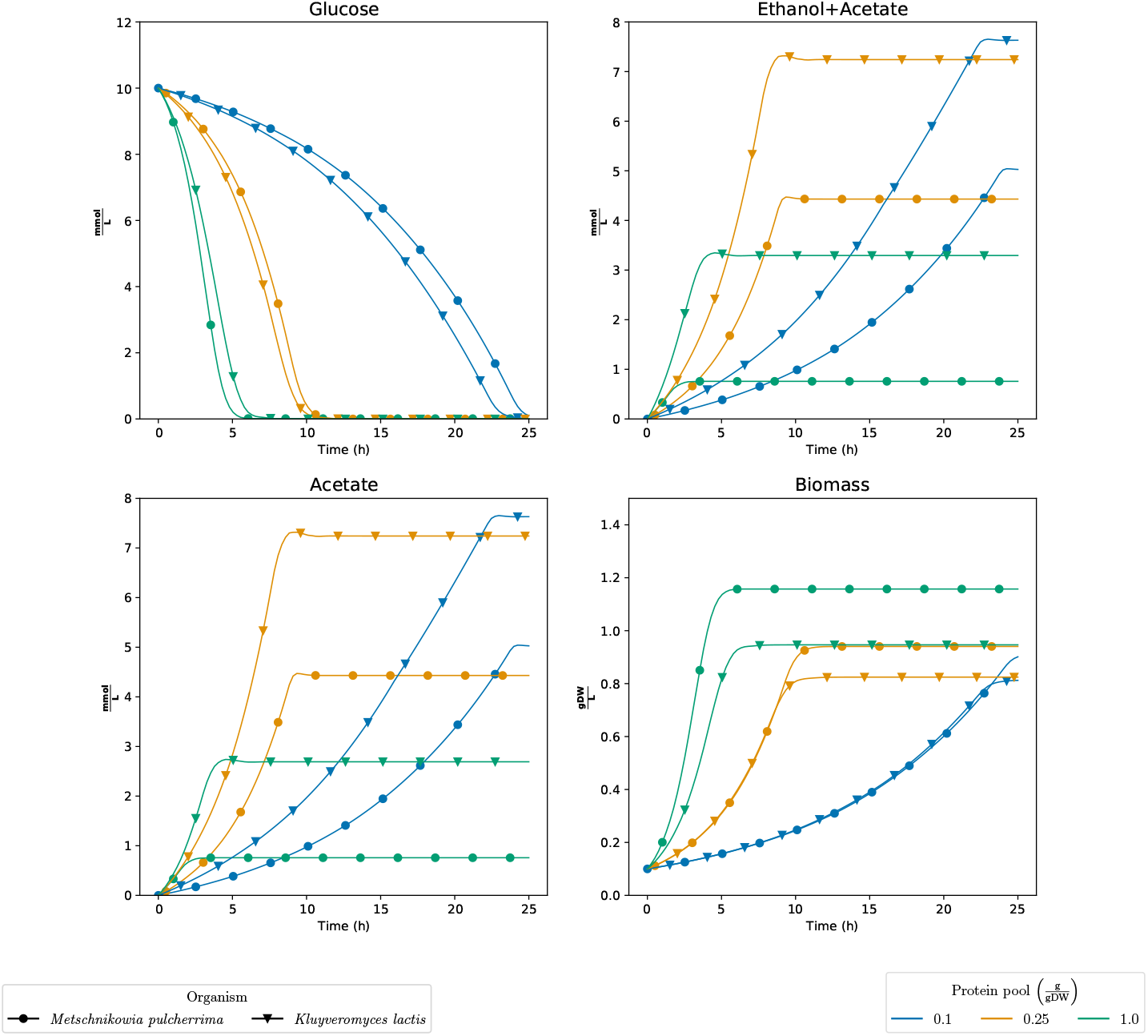
decFBA simulations of the sMOMENT models for *Metschnikowia pulcherrima* and *Kluyveromyces lactis* under different levels of the protein pool. 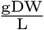 Grams of dry weight per liter.

## Discussion

In our metabolic model reconstructions, we found the GEM of *M. pulcherrima* to provide quantitatively different phenotypes compared to the other models, as it utilized glucose more efficiently, had a higher biomass yield, and conducted less fermentation. To a great extent, this echoes recent research which suggest *M. pulcherrima* as a good candidate for reducing alcohol content in wine[36, 16, 19, 22, 21, 46]. The connection between the three observed effects are quite straight forward. Respiration, instead of fermentation, gives better energy utilization of the substrate, less fermentation products and better growth for the same amount of substrate consumed.

Nonetheless, for certain yeast species, possessing an extensive respiratory metabolism may not confer an evolutionary advantage for two primary reasons. Firstly, the capacity to respire may not be advantageous in environments where oxygen supply is insufficient to sustain full respiration, and the production of ethanol inhibits competitors, as observed in commercial wine fermentation tanks[47, 41]. Secondly, in conditions characterized by high glucose concentrations, yeast may achieve elevated ATP production flux through ethanol fermentation as opposed to respiration, given that the latter necessitates greater protein utilization than the former[26, 25, 32, 48].

Our results suggest that the presumed presence of Complex I in *M. pulcherrima* allows the organism to respire glucose more efficiently than the other yeast strains. Thus, *M. pulcherrima* may be better adopted for respiration and will, therefore, prefer this mode of metabolism. Likewise, Malina *et al*.[25] attributed the high biomass yield and low production of ethanol and acetate of *Kluyveromyces marxianus* compared to other yeast strains, to be due to its presence of Complex I. At least for some obligate aerobic yeasts, such as *Yarrowia lipolytica, Rhodotorula muciluginosa*, and *Candida silvae*, Complex I is present, and inhibiting its activity[49, 42] leads to a reduction in respiration. According to Büschges *et al*., the yeasts with Complex I were suspected to use alternative NADH dehydrogenases when Complex I was inhibited, albeit with a penalty in growth, just as we observed with *M. pulcherrima*.

The sMOMENT models had shortcomings related to the enzyme constraints. First, carveFungi included reactions which were not annotated with a specific gene, meaning that these reactions in the downstream sMOMENT models, did not draw from the protein pool but were available without a protein cost. In particular, the mitochondrial genome was not sequenced for the non-*Saccharomyces* strains, meaning that the protein cost of some electron transport chain reactions were far from realistic values. Second, several metabolic enzymes exist as multimers, yet we did not have any information on the subunit stoichiometry. As a result, AutoPACMEN considered that exactly one of each distinct subunit was present in each complex. Furthermore, database *k*_*cat*_ values have been shown to be variable and often far from realistic *in vivo* values[50, 51, 52]. Finally, the *k*_*cat*_ coverage for non-model organisms is low, such that *S. cerevisiae* was likely the closest available candidate for picking *k*_*cat*_ values for many of the enzymes. We consider novel approaches for inferring *in vivo k*_*cat*_ from large-scale experimental data to be the best option for parameterizing high-quality models, although producing the data will be expensive an labour-intensive[51, 50, 53].

Our choice of parameters for dFBA simulations were based on educated guesses in lack of good data for calibration. Sànchez *et al*. used a enzymatic protein pool of *P*_*tot*_ = 0.448 g*/*gDW and saturation factor of *σ* = 0.5 for their ecYeast7 model of *S. cerevisiae*. In our case, this would correspond to a simulated protein pool of approximately 0.22 g*/*gDW since we assumed full saturation. Still, the protein cost of some enzymatic reactions were not accounted for in the sMOMENT models, so we think a somewhat lower enzyme pool would make a fairer comparison. Glucose uptake has been shown to vary considerably between different species of yeast and even between different strains of *S. cerevisiae*[54, 55]. From the available data and literature[56], we consider our chosen parameters to be within a realistic range.

Nevertheless, we acknowledge that glucose uptake and its balance to oxygen uptake is crucial to the nature of the fermentation. Less oxygen available compared to the consumption of glucose will favour fermentation at the expense of respiration. Additionally, regulatory mechanisms not accounted for by our models most likely also regulate the switching between fermentation and respiration[57]. Comparing Figure 1 and Supplementary Figure S1, we observed that the glucose concentration has a large effect on the production of fermented compounds, yet *M. pulcherrima* still has a stronger respiratory metabolism than the other yeasts for high glucose concentrations. We did not account for the fact that supplying oxygen is harder when the biomass concentration is high, making a fixed oxygen uptake of 10 mmol*/*gDW realistic in Figure 1, but unrealistic in Supplementary Figure S1. The model predictions for *M. pulcherrima* should inspire to further research and investigations into the industrial applications of respiratory yeasts. One of the central questions is whether our claim that *M. pulcherrima* has Complex I is correct, and if so, which phenotypic effects this enzyme has. Rotenone is known to be an inhibitor of Complex I and would therefore be a useful tool to study the activity of Complex I[42, 58, 59]. Systematic studies must be conducted in order to assess how *M. pulcherrima* behaves under varying availability of glucose and oxygen.

## Methods

### Creation of the yeast models

The protein sequences of the five species was obtained from the NCBI database[60, 61, 62, 63, 64, 65]. We annotated the function of the proteins with EggNog mapper V2[66] using Diamond[67] for the search of homologs in the EggNOG ortholog database version 5.

For the automatic model reconstruction, we used the software package CarveFungi[37]. CarveFungi is based on the CarveMe algorithm[33]. CarveFungi creates a score for each reaction in a universal metabolic model by linking their EC numbers to the annotation of the proteins obtained by EggNOG. The software contains a deep learning model to predict the subcellular localization of fungal proteins. This prediction contributes to the reaction score, assigning the reactions to a specific compartment in the model. The reaction scores are then used by a Mixed-Integer Linear Programming problem (MILP) to maximize the reactions present in the universal model with a high score and to minimize the reactions with a low score while maintaining the network connectivity and the model functionality.

The universal metabolic model employed for the reconstruction process was developed by integrating fungal reactions obtained from public databases such as KEGG[68] and MetaCyc[69]. This model was subsequently subjected to manual curation using relevant literature to ensure atom balance and simulatability, accomplished by incorporating exchange reactions and extending the biomass reaction based on the yeast consensus model[30].

The automated metabolic model reconstruction generated ensembles comprising up to 25 alternative models, each derived from the same genome. For the purpose of our analysis, we consolidated each ensemble into a single consensus model by including a reaction if it appeared in at least half of the models within the ensemble.

### Incorporation of enzymatic constraints

sMOMENT models with enzyme constraints were generated by feeding the GEMs into AutoPACMEN[34] version 0.6.1, applying default parameters. The BiGG metabolite file used by AutoPACMEN was retrieved from the BiGG[70] website (http://bigg.ucsd.edu/data_access, October 2020), while the BRENDA data was downloaded from the BRENDA[71] website (https://www.brenda-enzymes.org/download.php, October 2020). Before providing the models to AutoPACMEN, the models are augmented by Uniprot identifiers using Uniprot’s API. AutoPACMEN retrieved *k*_*cat*_ values from SABIO-RK[72, 73] and protein masses from Uniprot[44] using its built-in API interface (October 2020). AutoPACMEN’s model calibrator was not used.

### dFBA and decFBA simulations

The models of the five non-*Saccharomyces* strains and the iND750 *S. cerevisiae* model[39] were simulated *in silico* with dynamic FBA (dFBA)[38, 26]. The CO-BRApy package (version 0.25.0)[74] was used to handle the models, and the resulting LP problems were solved by the Gurobi optimizer (version 9.1.2). Glucose was the sole carbon source available with a maximum uptake flux determined by Michaelis-Menten kinetics: 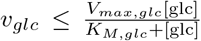, where the maximal up-take rate *V*_*max,glc*_ = 10 mmol*/*gDW, the half-saturation constant *K*_*M,glc*_ = 5 mmol, and [glc] was the glucose concentration in the medium which was initiated to [glc]_0_ = 10 mmol L^−1^ for all simulations expect for Supplementary Figure S1 where [glc]_0_ = 1000 mmol L^−1^. The biomass concentration was as initiated to [X]_0_ = 0.1 gDW*/*L. Oxygen was available at a fixed rate of *v*_oxygen_ ≤ 10 mmol*/*gDW. We monitored several entities, including biomass, glucose, acetate, ethanol, and glycerol. The latter three components were incorporated to track the accumulation of fermentation products generated by the yeast. However, under the tested conditions, none of the models produced glycerol; hence, we omitted its depiction for clarity. Furthermore, we sought to highlight the combined production of ethanol and acetate. Consequently, the graphical representations included a panel displaying the sum of ethanol and acetate concentrations in the medium, and another panel illustrating the acetate concentration separately. To preclude physiologically implausible metabolic exports, we blocked the export reactions for lactate (both stereoisomers), dihydroxyacetate, D-ribulose, arabinitol, and ribose. Additionally, we inhibited a wasteful mitochondrial membrane proton leakage reaction, which would have led to physiologically unreasonable outcomes if not removed from the model.

To obtain consistent and physiologically plausible results, we applied lexicographic objectives when performing FBA on the models prioritized in the following order:

1. Maximize production of biomass
2. Minimize consumption of glucose
3. Maximize excretion of ethanol
4. Maximize excretion of acetate
5. Maximize excretion of glycerol

The models were simulated using the static optimization approach and SciPy’s `solve_ivp` function[75]. For the ODE solver, the BDF algorithm[76] was used with an absolute and relative tolerance of 10^−2^. In cases where the optimization problem became infeasible, the simulation was terminated, but results were padded such that the final state of the system was imputed to all time-points beyond the termination. This happened only if the model was unable to grow because the carbon source (glucose) in the medium was depleted.

dFBA was performed both for the original models generated with CarveFungi and the sMOMENT models processed through AutoPACMEN. Upon running decFBA with the sMOMENT models, the level of the enzyme pool was adjusted. Three different levels of the enzymatic protein pool (0.1, 0.25, and 1.0 grams of protein per gram dry weight(g/gWD)) were chosen.

## Abbreviations

OIV: International Organisation of Vine and Wine
FBA: Flux Balance Analysis
dFBA: Dynamic Flux Balance Analysis
decFBA: Dynamic Enzyme-Constrained FBA
MILP: Mixed-Integer Linear Programming
GEM: Genome-Scale metabolic Model
ecGEM: Enzyme Constrained Genome-Scale metabolic Model
LP: Linear Programming
DW: Dry Weight
PMF: Proton Motive Force
ODE: Ordinary Differential Equation

## Declarations

### Ethics approval and consent to participate

Not applicable.

### Consent for publication

Not applicable.

### Availability of data and material

The source code and data for generating, augmenting and analysing the GEMs used in this paper in addition to the resulting GEMs is available at Figshare (http://dx.doi.org/10.6084/m9.figshare.22664848).

### Authors’ contributions

S.C. and P.J. created the GEMs from the genome sequences using CarveFungi and conducted the bioinformatics analyses. J.P.P. augmented the GEMs with enzyme constraints using AutoPACMEN, ran simulations, studied the properties of the GEMs, and wrote the first draft of the paper. E.A. supervised the project. All authors contributed to and accepted the final version of the paper.

## Acknowledgements

Thanks to Pavlos Stephanos Bekiaris for providing technical support and assistance with the AutoPACMEN software. Thanks to Ramon Gonzalez for input on the project and for reading and providing comments on the manuscript.

## Author details

^1^Department of Biotechnology and Food Science, NTNU-Norwegian University of Science and Technology, Trondheim, Norway. ^2^K.G. Jebsen Center for Genetic Epidemiology, Department of Public Health and General Practice, NTNU-Norwegian University of Science and Technology, Trondheim, Trondheim, Norway. ^3^VTT Technical Research Centre of Finland, Espoo, Finland. ^4^Department of Bioproducts and Biosystems Aalto University, Espoo, Finland.

## Additional Files

Additional file 1

Supplementary figures

## References

1. Lin, X., Tang, X., Han, X., He, X., Han, N., Ding, Y., Sun, Y.: Effect of metschnikowia pulcherrima on saccharomyces cerevisiae pdh by-pass in mixedfermentation with varied sugar concentrations of synthetic grape juice and inoculation ratios. Fermentation 8(10) (2022). doi:10.3390/fermentation8100480

2. Contreras, A., Curtin, C., Varela, C.: Yeast population dynamics reveal a potential ‘collaboration’ between metschnikowia pulcherrima and saccharomyces uvarum for the production of reduced alcohol wines during shiraz fermentation. Applied Microbiology and Biotechnology 99(4), 1885–1895 (2015). doi:10.1007/s00253-014-6193-6

3. García, M., Esteve-Zarzoso, B., Cabellos, J.M., Arroyo, T.: Sequential non-saccharomyces and saccharomyces cerevisiae fermentations to reduce the alcohol content in wine. Fermentation 6(2) (2020). doi:10.3390/fermentation6020060

4. Jolly, N.P., Varela, C., Pretorius, I.S.: Not your ordinary yeast: non-Saccharomyces yeasts in wine production uncovered. FEMS Yeast Research 14(2), 215–237 (2014). doi:10.1111/1567-1364.12111

5. Ciani, M., Morales, P., Comitini, F., Tronchoni, J., Canonico, L., Curiel, J.A., Oro, L., Rodrigues, A.J., Gonzalez, R.: Non-conventional yeast species for lowering ethanol content of wines. Frontiers in Microbiology 7 (2016). doi:10.3389/fmicb.2016.00642

6. Gonzalez, R., Quirós, M., Morales, P.: Yeast respiration of sugars by non-saccharomyces yeast species: A promising and barely explored approach to lowering alcohol content of wines. Trends in Food Science & Technology 29(1), 55–61 (2013). doi:10.1016/j.tifs.2012.06.015

7. Tufariello, M., Fragasso, M., Pico, J., Panighel, A., Castellarin, S.D., Flamini, R., Grieco, F.: Influence of non-saccharomyces on wine chemistry: A focus on aroma-related compounds. Molecules 26(3), 644 (2021). doi:10.3390/molecules26030644

8. Varela, C., Dry, P.R., Kutyna, D.R., Francis, I.L., Henschke, P.A., Curtin, C.D., Chambers, P.J.: Strategies for reducing alcohol concentration in wine. Australian Journal of Grape and Wine Research 21(S1), 670–679 (2015). doi:10.1111/ajgw.12187

9. Alston, J., Fuller, K.B., Lapsley, J.T., Soleas, G.: Too much of a good thing? causes and consequences of increases in sugar content of california wine grapes*. Journal of Wine Economics 6(2), 135–159 (2011)

10. Longo, R., Blackman, J.W., Torley, P.J., Rogiers, S.Y., Schmidtke, L.M.: Changes in volatile composition and sensory attributes of wines during alcohol content reduction. Journal of the Science of Food and Agriculture 97(1), 8–16 (2017). doi:10.1002/jsfa.7757

11. International Organisation of Vine and Wine: International oenological codex. Regulatory document, International Organisation of Vine and Wine (2023). https://www.oiv.int/sites/default/files/publication/2023-04/CODEXcomplet2023EN.pdf

12. The European Commission: Commission delegated regulation (eu) 2019/934 of 12 march 2019 supplementing regulation (eu) no 1308/2013 of the european parliament and of the council as regards wine-growing areas where the alcoholic strength may be increased, authorised oenological practices and restrictions applicable to the production and conservation of grapevine products, the minimum percentage of alcohol for by-products and their disposal, and publication of oiv files. Regulatory document, The European Commission (2019). https://eur-lex.europa.eu/legal-content/EN/TXT/PDF/?uri=CELEX:32019R0934

13. Bärwald, G., Fischer, A.: Crabtree effect in aerobic fermentations using grape juice for the production of alcohol reduced wine. Biotechnology Letters 18(10), 1187–1192 (1996). doi:10.1007/BF00128590

14. Hammad, N., Rosas-Lemus, M., Uribe-Carvajal, S., Rigoulet, M., Devin, A.: The crabtree and warburg effects: Do metabolite-induced regulations participate in their induction? Biochimica et biophysica acta 1857, 1139–1146 (2016). doi:10.1016/j.bbabio.2016.03.034

15. Curiel, J.A., Salvadó, Z., Tronchoni, J., Morales, P., Rodrigues, A.J., Quirós, M., Gonzalez, R.: Identification of target genes to control acetate yield during aerobic fermentation with saccharomyces cerevisiae. Microbial Cell Factories 15(1), 156 (2016). doi:10.1186/s12934-016-0555-y

16. Morales, P., Rojas, V., Quirós, M., Gonzalez, R.: The impact of oxygen on the final alcohol content of wine fermented by a mixed starter culture. Applied Microbiology and Biotechnology 99(9), 3993–4003 (2015). doi:10.1007/s00253-014-6321-3

17. van Dijken, J.P., Weusthuis, R.A., Pronk, J.T.: Kinetics of growth and sugar consumption in yeasts. Antonie van Leeuwenhoek 63, 343–352 (1993). doi:10.1007/BF00871229

18. Vicente, J., Ruiz, J., Belda, I., Benito-Vázquez, I., Marquina, D., Calderón, F., Santos, A., Benito, S.: The genus metschnikowia in enology. Microorganisms 8(7) (2020). doi:10.3390/microorganisms8071038

19. Tronchoni, J., Curiel, J.A., Sáenz-Navajas, M.P., Morales, P., de-la-Fuente-Blanco, A., Fernández-Zurbano, P., Ferreira, V., Gonzalez, R.: Aroma profiling of an aerated fermentation of natural grape must with selected yeast strains at pilot scale. Food microbiology 70, 214–223 (2018). doi:10.1016/j.fm.2017.10.008

20. Canonico, L., Comitini, F., Oro, L., Ciani, M.: Sequential fermentation with selected immobilized non-saccharomyces yeast for reduction of ethanol content in wine. Frontiers in Microbiology 7 (2016). doi:10.3389/fmicb.2016.00278

21. Hranilovic, A., Gambetta, J.M., Jeffery, D.W., Grbin, P.R., Jiranek, V.: Lower-alcohol wines produced by metschnikowia pulcherrima and saccharomyces cerevisiae co-fermentations: The effect of sequential inoculation timing. International Journal of Food Microbiology 329, 108651 (2020). doi:10.1016/j.ijfoodmicro.2020.108651

22. Contreras, A., Hidalgo, C., Henschke, P.A., Chambers, P.J., Curtin, C., Varela, C.: Evaluation of non-saccharomyces yeasts for the reduction of alcohol content in wine. Applied and environmental microbiology 80, 1670–8 (2014). doi:10.1128/AEM.03780-13

23. Passi, A., Tibocha-Bonilla, J.D., Kumar, M., Tec-Campos, D., Zengler, K., Zuniga, C.: Genome-scale metabolic modeling enables in-depth understanding of big data. Metabolites 12 (2021)

24. Gu, C., Kim, G.B., Kim, W.J., Kim, H.U., Lee, S.Y.: Current status and applications of genome-scale metabolic models. Genome Biology 20(1), 121 (2019). doi:10.1186/s13059-019-1730-3

25. Malina, C., Yu, R., Bj rkeroth, J., Kerkhoven, E.J., Nielsen, J.: Adaptations in metabolism and protein translation give rise to the crabtree effect in yeast. Proceedings of the National Academy of Sciences 118(51), 2112836118 (2021). doi:10.1073/pnas.2112836118

26. Moreno-Paz, S., Schmitz, J., Martins Dos Santos, V.A.P., Suarez-Diez, M.: Enzyme-constrained models predict the dynamics of saccharomyces cerevisiae growth in continuous, batch and fed-batch bioreactors. Microbial biotechnology (2022). doi:10.1111/1751-7915.13995

27. Jouhten, P., Konstantinidis, D., Pereira, F., Andrejev, S., Grkovska, K., Castillo, S., Ghiachi, P., Beltran, G., Almaas, E., Mas, A., Warringer, J., Gonzalez, R., Morales, P., Patil, K.R.: Predictive evolution of metabolic phenotypes using model-designed environments. Molecular Systems Biology 18(10), 10980 (2022). doi:10.15252/msb.202210980

28. Scott, W.T., Smid, E.J., Notebaart, R.A., Block, D.E.: Curation and analysis of a saccharomyces cerevisiae genome-scale metabolic model for predicting production of sensory impact molecules under enological conditions. Processes 8(9) (2020). doi:10.3390/pr8091195

29. Henriques, D., Minebois, R., Mendoza, S.N., Macías, L.G., Pérez-Torrado, R., Barrio, E., Teusink, B., Querol, A., Balsa-Canto, E., Vecchio, D.D.: A multiphase multiobjective dynamic genome-scale model shows different redox balancing among yeast species of the saccharomyces genus in fermentation. mSystems 6(4), 00260–21 (2021). doi:10.1128/mSystems.00260-21

30. Lu, H., Li, F., Sánchez, B.J., Zhu, Z., Li, G., Domenzain, I., Marcisauskas, S., Anton, P.M., Lappa, D., Lieven, C., Beber, M.E., Sonnenschein, N., Kerkhoven, E.J., Nielsen, J.: A consensus s. cerevisiae metabolic model yeast8 and its ecosystem for comprehensively probing cellular metabolism. Nature Communications 10(1), 3586 (2019). doi:10.1038/s41467-019-11581-3

31. Lu, H., Li, F., Yuan, L., Domenzain, I., Yu, R., Wang, H., Li, G., Chen, Y., Ji, B., Kerkhoven, E.J., Nielsen, J.: Yeast metabolic innovations emerged via expanded metabolic network and gene positive selection. Molecular Systems Biology 17(10), 10427 (2021). doi:10.15252/msb.202110427

32. Nilsson, A., Nielsen, J.: Metabolic trade-offs in yeast are caused by f1f0-atp synthase. Scientific Reports 6(1), 22264 (2016). doi:10.1038/srep22264

33. Machado, D., Andrejev, S., Tramontano, M., Patil, K.R.: Fast automated reconstruction of genome-scale metabolic models for microbial species and communities. Nucleic Acids Research 46(15), 7542–7553 (2018). doi:10.1093/nar/gky537

34. Bekiaris, P.S., Klamt, S.: Automatic construction of metabolic models with enzyme constraints. BMC Bioinformatics 21(1), 19 (2020). doi:10.1186/s12859-019-3329-9

35. Sánchez, B.J., Zhang, C., Nilsson, A., Lahtvee, P.-J., Kerkhoven, E.J., Nielsen, J.: Improving the phenotype predictions of a yeast genome-scale metabolic model by incorporating enzymatic constraints. Mol Syst Biol 13(8), 935 (2017). doi:10.15252/msb.20167411

36. Quirós, M., Rojas, V., Gonzalez, R., Morales, P.: Selection of non-saccharomyces yeast strains for reducing alcohol levels in wine by sugar respiration. International journal of food microbiology 181, 85–91 (2014). doi:10.1016/j.ijfoodmicro.2014.04.024

37. Castillo, S.: CarveFungi (2021). https://github.com/SandraCastilloPriego/CarveFungi

38. Mahadevan, R., Edwards, J.S., Doyle, F.J. 3rd: Dynamic flux balance analysis of diauxic growth in escherichia coli. Biophys J 83(3), 1331–40 (2002). doi:10.1016/S0006-3495(02)73903-9

39. Duarte, N.C., Herrgård, M.J., Palsson, B.o.: Reconstruction and validation of saccharomyces cerevisiae ind750, a fully compartmentalized genome-scale metabolic model. Genome research 14, 1298–309 (2004). doi:10.1101/gr.2250904

40. Antos-Krzeminska, N., Jarmuszkiewicz, W.: Alternative type ii nad(p)h dehydrogenases in the mitochondria of protists and fungi. Protist 170(1), 21–37 (2019). doi:10.1016/j.protis.2018.11.001

41. Hagman, A., Säll, T., Compagno, C., Piskur, J.: Yeast “make-accumulate-consume” life strategy evolved as a multi-step process that predates the whole genome duplication. PLOS ONE 8(7), 1–12 (2013). doi:10.1371/journal.pone.0068734

42. Büschges, R., Bahrenberg, G., Zimmermann, M., Wolf, K.: Nadh: Ubiquinone oxidoreductase in obligate aerobic yeasts. Yeast 10(4), 475–479 (1994). doi:10.1002/yea.320100406

43. Altschul, S.F., Gish, W., Miller, W., Myers, E.W., Lipman, D.J.: Basic local alignment search tool. Journal of molecular biology 215, 403–10 (1990). doi:10.1016/S0022-2836(05)80360-2

44. Bateman, A., Martin, M.-J., Orchard, S., Magrane, M., Agivetova, R., Ahmad, S., Alpi, E., Bowler-Barnett, E.H., Britto, R., Bursteinas, B., Bye-A-Jee, H., Coetzee, R., Cukura, A., Da Silva, A., Denny, P., Dogan, T., Ebenezer, T., Fan, J., Castro, L.G., Garmiri, P., Georghiou, G., Gonzales, L., Hatton-Ellis, E., Hussein, A., Ignatchenko, A., Insana, G., Ishtiaq, R., Jokinen, P., Joshi, V., Jyothi, D., Lock, A., Lopez, R., Luciani, A., Luo, J., Lussi, Y., Mac-Dougall, A., Madeira, F., Mahmoudy, M., Menchi, M., Mishra, A., Moulang, K., Nightingale, A., Oliveira, C.S., Pundir, S., Qi, G., Raj, S., Rice, D., Lopez, M.R., Saidi, R., Sampson, J., Sawford, T., Speretta, E., Turner, E., Tyagi, N., Vasudev, P., Volynkin, V., Warner, K., Watkins, X., Zaru, R., Zellner, H., Bridge, A., Poux, S., Redaschi, N., Aimo, L., Argoud-Puy, G., Auchincloss, A., Axelsen, K., Bansal, P., Baratin, D., Blatter, M.-C., Bolleman, J., Boutet, E., Breuza, L., Casals-Casas, C., de Castro, E., Echioukh, K.C., Coudert, E., Cuche, B., Doche, M., Dornevil, D., Estreicher, A., Famiglietti, M.L., Feuermann, M., Gasteiger, E., Gehant, S., Gerritsen, V., Gos, A., Gruaz-Gumowski, N., Hinz, U., Hulo, C., Hyka-Nouspikel, N., Jungo, F., Keller, G., Kerhornou, A., Lara, V., Le Mercier, P., Lieberherr, D., Lombardot, T., Martin, X., Masson, P., Morgat, A., Neto, T.B., Paesano, S., Pedruzzi, I., Pilbout, S., Pourcel, L., Pozzato, M., Pruess, M., Rivoire, C., Sigrist, C., Sonesson, K., Stutz, A., Sundaram, S., Tognolli, M., Verbregue, L., Wu, C.H., Arighi, C.N., Arminski, L., Chen, C., Chen, Y., Garavelli, J.S., Huang, H., Laiho, K., McGarvey, P., Natale, D.A., Ross, K., Vinayaka, C.R., Wang, Q., Wang, Y., Yeh, L.-S., Zhang, J., Consortium, U.: Uniprot: the universal protein knowledgebase in 2021. NUCLEIC ACIDS RESEARCH 49(D1), 480–489 (2021). doi:10.1093/nar/gkaa1100

45. Karlsen, E., Gylseth, M., Schulz, C., Almaas, E.: A study of a diauxic growth experiment using an expanded dynamic flux balance framework. PLOS ONE 18(1), 1–17 (2023). doi:10.1371/journal.pone.0280077

46. Canonico, L., Comitini, F., Ciani, M.: Metschnikowia pulcherrima selected strain for ethanol reduction in wine: Influence of cell immobilization and aeration condition. Foods (Basel, Switzerland) 8 (2019). doi:10.3390/foods8090378

47. Hagman, A., Piskur, J.: A study on the fundamental mechanism and the evolutionary driving forces behind aerobic fermentation in yeast. PLOS ONE 10(1), 1–24 (2015). doi:10.1371/journal.pone.0116942

48. Pfeiffer, T., Morley, A.: An evolutionary perspective on the crabtree effect. Frontiers in Molecular Biosciences 1 (2014). doi:10.3389/fmolb.2014.00017

49. Bych, K., Kerscher, S., Netz, D.J.A., Pierik, A.J., Zwicker, K., Huynen, M.A., Lill, R., Brandt, U., Balk, J.: The iron–sulphur protein ind1 is required for effective complex i assembly. The EMBO Journal 27(12), 1736–1746 (2008). doi:10.1038/emboj.2008.98

50. Heckmann, D., Campeau, A., Lloyd, C.J., Phaneuf, P.V., Hefner, Y., Carrillo-Terrazas, M., Feist, A.M., Gonzalez, D.J., Palsson, B.O.: Kinetic profiling of metabolic specialists demonstrates stability and consistency of in vivo enzyme turnover numbers. Proceedings of the National Academy of Sciences of the United States of America 117, 23182–23190 (2020). doi:10.1073/pnas.2001562117

51. Heckmann, D., Lloyd, C.J., Mih, N., Ha, Y., Zielinski, D.C., Haiman, Z.B., Desouki, A.A., Lercher, M.J., Palsson, B.O.: Machine learning applied to enzyme turnover numbers reveals protein structural correlates and improves metabolic models. Nature Communications 9(1), 5252 (2018). doi:10.1038/s41467-018-07652-6

52. Wendering, P., Arend, M., Razaghi-Moghadamkashani, Z., Nikoloski, Z.: Data integration across conditions improves turnover number estimates and metabolic predictions. bioRxiv (2022). doi:10.1101/2022.04.01.486742

53. Li, F., Yuan, L., Lu, H., Li, G., Chen, Y., Engqvist, M.K.M., Kerkhoven, E.J., Nielsen, J.: Deep learning-based kcat prediction enables improved enzyme-constrained model reconstruction. Nature Catalysis 5(8), 662–672 (2022). doi:10.1038/s41929-022-00798-z

54. Does, A.L., Bisson, L.F.: Comparison of glucose uptake kinetics in different yeasts. Journal of bacteriology 171, 1303–8 (1989)

55. Nissen, P., Nielsen, D., Arneborg, N.: The relative glucose uptake abilities of non-saccharomyces yeasts play a role in their coexistence with saccharomyces cerevisiae in mixed cultures. Applied Microbiology and Biotechnology 64(4), 543–550 (2004). doi:10.1007/s00253-003-1487-0

56. Pizarro, F., Varela, C., Martabit, C., Bruno, C., Pérez-Correa, J.R., Agosin, E.: Coupling kinetic expressions and metabolic networks for predicting wine fermentations. Biotechnology and Bioengineering 98(5), 986–998 (2007). doi:10.1002/bit.21494

57. Otterstedt, K., Larsson, C., Bill, R.M., Ståhlberg, A., Boles, E., Hohmann, S., Gustafsson, L.: Switching the mode of metabolism in the yeast saccharomyces cerevisiae. EMBO reports 5(5), 532–537 (2004). doi:10.1038/sj.embor.7400132

58. Heinz, S., Freyberger, A., Lawrenz, B., Schladt, L., Schmuck, G., Ellinger-Ziegelbauer, H.: Mechanistic investigations of the mitochondrial complex i inhibitor rotenone in the context of pharmacological and safety evaluation. Scientific Reports 7(1), 45465 (2017). doi:10.1038/srep45465

59. Ozay, E.I., Sherman, H.L., Mello, V., Trombley, G., Lerman, A., Tew, G.N., Yadava, N., Minter, L.M.: Rotenone treatment reveals a role for electron transport complex i in the subcellular localization of key transcriptional regulators during t helper cell differentiation. Frontiers in immunology 9, 1284 (2018). doi:10.3389/fimmu.2018.01284

60. O’Leary, N.A., Wright, M.W., Brister, J.R., Ciufo, S., Haddad, D., McVeigh, R., Rajput, B., Robbertse, B., Smith-White, B., Ako-Adjei, D., Astashyn, A., Badretdin, A., Bao, Y., Blinkova, O., Brover, V., Chetvernin, V., Choi, J., Cox, E., Ermolaeva, O., Farrell, C.M., Goldfarb, T., Gupta, T., Haft, D., Hatcher, E., Hlavina, W., Joardar, V.S., Kodali, V.K., Li, W., Maglott, D., Masterson, P., McGarvey, K.M., Murphy, M.R., O’Neill, K., Pujar, S., Rangwala, S.H., Rausch, D., Riddick, L.D., Schoch, C., Shkeda, A., Storz, S.S., Sun, H., Thibaud-Nissen, F., Tolstoy, I., Tully, R.E., Vatsan, A.R., Wallin, C., Webb, D., Wu, W., Landrum, M.J., Kimchi, A., Tatusova, T., DiCuccio, M., Kitts, P., Murphy, T.D., Pruitt, K.D.: Reference sequence (RefSeq) database at NCBI: current status, taxonomic expansion, and functional annotation 44, 733–745. doi:10.1093/nar/gkv1189. Accessed 2022-12-11

61. Kluyveromyces Lactis, Assembly ASM251v1. National Center for Biotechnology Information (NCBI). https://www.ncbi.nlm.nih.gov/genome/?term=ASM251v1

62. Metschnikowia Pulcherrima, Assembly ASM421770v1. National Center for Biotechnology Information (NCBI). https://www.ncbi.nlm.nih.gov/assembly/GCA_004217705.1/

63. Lachancea Thermotolerans, Assembly ASM14280v1. National Center for Biotechnology Information (NCBI). https://www.ncbi.nlm.nih.gov/genome/?term=ASM14280v1

64. Torulaspora Delbrueckii, Assembly ASM24337v1. National Center for Biotechnology Information (NCBI). https://www.ncbi.nlm.nih.gov/data-hub/genome/GCF_000243375.1/

65. Hanseniaspora Osmophila, Assembly ASM174704v1. National Center for Biotechnology Information (NCBI). https://www.ncbi.nlm.nih.gov/genome/46405?genome_assembly_id=283698

66. Huerta-Cepas, J., Szklarczyk, D., Heller, D., Hernández-Plaza, A., Forslund, S.K., Cook, H., Mende, D.R., Letunic, I., Rattei, T., Jensen, L.J., von Mering, C., Bork, P.: eggNOG 5.0: a hierarchical, functionally and phylogenetically annotated orthology resource based on 5090 organisms and 2502 viruses 47, 309–314. doi:10.1093/nar/gky1085

67. Buchfink, B., Reuter, K., Drost, H.-G.: Sensitive protein alignments at tree-of-life scale using diamond. Nature Methods 18(4), 366–368 (2021). doi:10.1038/s41592-021-01101-x

68. Kanehisa, M., Goto, S.: KEGG: kyoto encyclopedia of genes and genomes 28(1), 27–30. doi:10.1093/nar/28.1.27

69. Caspi, R., Billington, R., Ferrer, L., Foerster, H., Fulcher, C.A., Keseler, I.M., Kothari, A., Krummenacker, M., Latendresse, M., Mueller, L.A., Ong, Q., Paley, S., Subhraveti, P., Weaver, D.S., Karp, P.D.: The MetaCyc database of metabolic pathways and enzymes and the BioCyc collection of pathway/genome databases 44, 471–480. doi:10.1093/nar/gkv1164

70. King, Z.A., Lu, J., Draeger, A., Miller, P., Federowicz, S., Lerman, J.A., Ebrahim, A., Palsson, B.O., Lewis, N.E.: Bigg models: A platform for integrating, standardizing and sharing genome-scale models. NUCLEIC ACIDS RESEARCH 44(D1), 515–522 (2016). doi:10.1093/nar/gkv1049

71. Chang, A., Jeske, L., Ulbrich, S., Hofmann, J., Koblitz, J., Schomburg, I., Neumann-Schaal, M., Jahn, D., Schomburg, D.: Brenda, the elixir core data resource in 2021: new developments and updates. NUCLEIC ACIDS RESEARCH 49(D1), 498–508 (2021). doi:10.1093/nar/gkaa1025

72. Wittig, U., Kania, R., Golebiewski, M., Rey, M., Shi, L., Jong, L., Algaa, E., Weidemann, A., Sauer-Danzwith, H., Mir, S., Krebs, O., Bittkowski, M., Wetsch, E., Rojas, I., Mueller, W.: Sabio-rk-database for biochemical reaction kinetics. NUCLEIC ACIDS RESEARCH 40(D1), 790–796 (2012). doi:10.1093/nar/gkr1046

73. Wittig, U., Rey, M., Weidemann, A., Kania, R., Mueller, W.: Sabio-rk: an updated resource for manually curated biochemical reaction kinetics. NUCLEIC ACIDS RESEARCH 46(D1), 656–660 (2018). doi:10.1093/nar/gkx1065

74. Ebrahim, A., Lerman, J.A., Palsson, B.O., Hyduke, D.R.: Cobrapy: Constraints-based reconstruction and analysis for python. BMC Systems Biology 7(1), 74 (2013). doi:10.1186/1752-0509-7-74

75. Virtanen, P., Gommers, R., Oliphant, T.E., Haberland, M., Reddy, T., Cournapeau, D., Burovski, E., Peterson, P., Weckesser, W., Bright, J., van der Walt, S.J., Brett, M., Wilson, J., Millman, K.J., Mayorov, N., Nelson, A.R.J., Jones, E., Kern, R., Larson, E., Carey, C.J., Polat, İ., Feng, Y., Moore, E.W., VanderPlas, J., Laxalde, D., Perktold, J., Cimrman, R., Henriksen, I., Quintero, E.A., Harris, C.R., Archibald, A.M., Ribeiro, A.H., Pedregosa, F., van Mulbregt, P., SciPy 1.0 Contributors: SciPy 1.0: Fundamental Algorithms for Scientific Computing in Python. Nature Methods 17, 261–272 (2020). doi:10.1038/s41592-019-0686-2

76. Byrne, G.D., Hindmarsh, A.C.: A polyalgorithm for the numerical solution of ordinary differential equations. ACM Trans. Math. Softw. 1(1), 71–96 (1975). doi:10.1145/355626.355636

