## Supplementary Materials for "Genome-scale metabolic models reveal determinants of phenotypic differences in non-Saccharomyces yeasts"

### Supplementary material for "Genome-scale metabolic models reveal determinants of phenotypic differences in non-Saccharomyces yeasts"

**Jakob Peder Pettersen, Sandra Castillo, Paula Jouhten, Eivind Almaas**

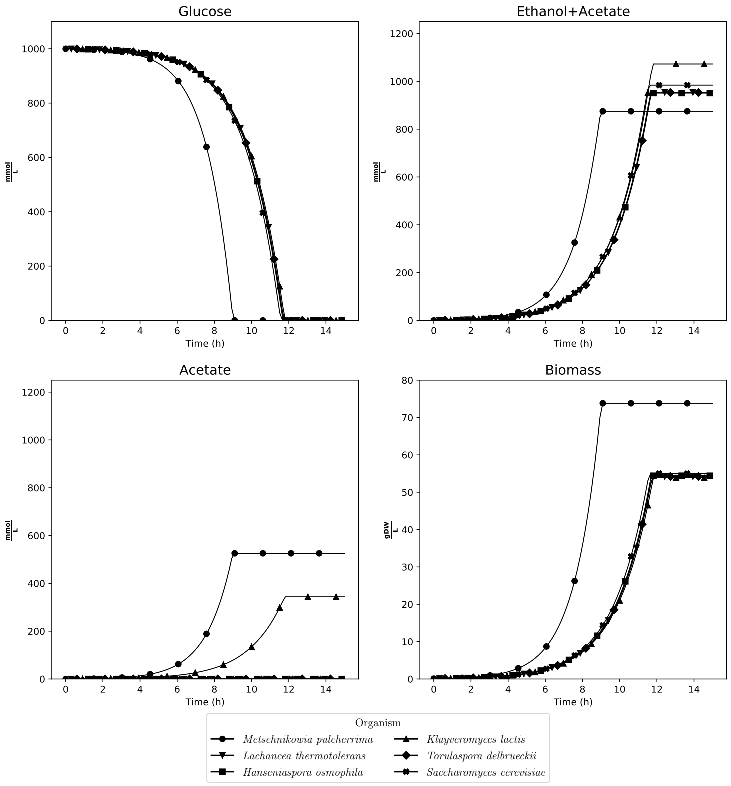


Figure S1: dFBA simulations of the models without enzymatic constraints for the six yeast strains, starting with $1000\,\mathrm{mmol\,L^{-1}}$ glucose. $\frac{\mathrm{gDW}}{\mathrm{L}}$: Grams of dry weight per liter.

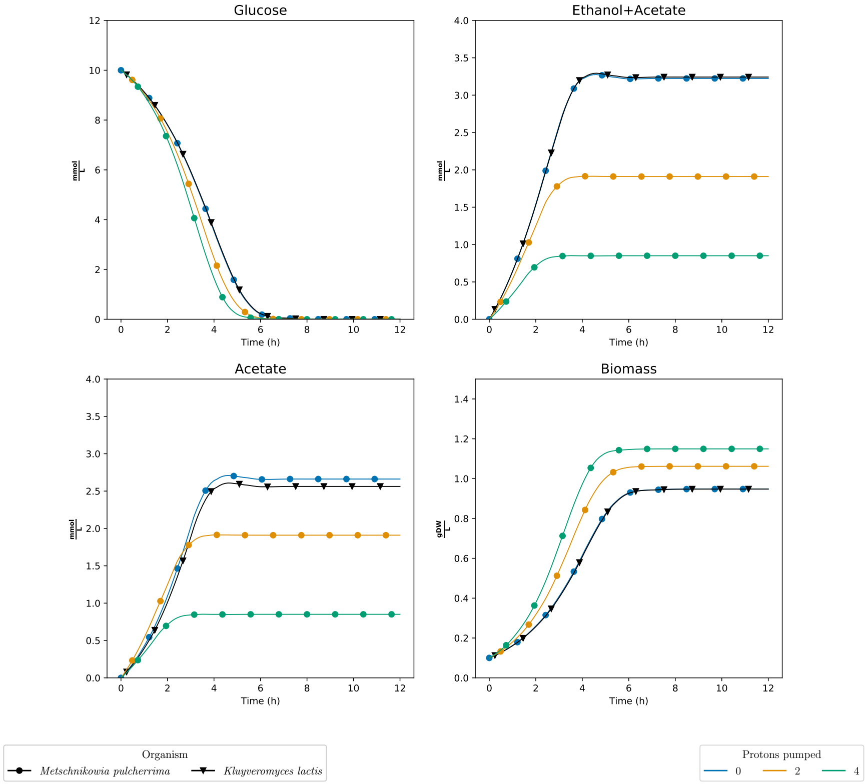


Figure S2: dFBA simulations of the models without enzymatic constraints in *Metschnikowia pulcherrima* and *Kluyveromyces lactis* when artificially changing the stoichiometry of the number of protons pumped by Complex I. In these simulations, the reactions L-glutamate:NADP+ oxidoreductase and Isocitrate:NADP+ oxidoreductase for *Metschnikowia pulcherrima* are knocked out.
